# Video observations of the acute effects of dredged sediment deposition on mobile epifauna

**DOI:** 10.1101/496281

**Authors:** G.C. Roegner, S.A. Fields, S.K. Henkel

## Abstract

The “beneficial uses” of dredged sediment are increasingly being explored for habitat restoration and beach nourishment. At ocean and nearshore deposition sites, any beneficial use must be tempered by evaluating impacts to the benthos. We studied a site at the mouth of the Columbia River where a “thin-layer” sediment deposition method was employed to minimize mounding and disperse sediment within a prescribed area. We used baited benthic video landers (BVLs) in a Before-After Control-Impact (BACI) experimental design to test the acute effects of sediment deposition on the Dungeness crab (*Cancer magister*) and dog whelk (*Nassarius* spp). We considered the acute effects of both sediment deposition depths and the lateral surge (the turbidity front transiting the seafloor). Observations revealed sedimentation levels were limited (< 4 cm) and likely posed no direct threat to epifauna. Video and instrument readings showed the lateral surge to impact the BVLs as a 2 to 3 m/s sediment-laden front. Crabs were significantly impacted, while gastropods were more resistant to dislodgment. However, the high velocity impact was relatively brief (5 to 7 min). Further, crabs often returned to forage at BVLs after a mean lag of about 20 min post-impact. These results indicate an acute but ephemeral impact effect on crab, and support use of the thin-layer deposition method to minimize burial. The BVLs in a BACI experimental design were an effective means of measuring sediment impacts to mobile epifauna, and video observations were informative for understanding lateral surge dynamics and the behavioral interactions of organisms.

## 1.0 Introduction

Recent advances in dredged sediment management include recognition of the potential to mitigate areas of environmental degradation with judicious sediment placement [1–3]. Such “beneficial uses” include creation, restoration, and maintenance of wetland habitat, and nourishment of eroding ocean beaches. In the Pacific Northwest of the USA, reduced riverine sediment supply [4] and increased wave heights [5] are among the leading causes of ocean beach erosion that has threatened coastal communities and infrastructure, including the ocean jetties stabilizing the mouth of the Columbia River (MCR). Beneficial sediment deposition in the nearshore zone has been proposed to mitigate for the shoreline recession at the MCR [6]. However, designated deposition areas are productive Dungeness crab (*Cancer magister*) fishing grounds. This has prompted concerns over the impacts of deposition events on crab and other fauna (as well as on navigation interests), and spurred research to quantify possible deleterious effects [7, 8]. Here we report experiments on the acute *in situ* effects of sediment deposition on the Dungeness crab, a dominant mobile epibenthic predator and important fishery resource.

Sediment deposition events can harm marine fauna through increased turbidity or toxic pollution levels [9–11], as well as from direct burial or mechanical effects of the sediment-laden plume as it impacts and transits the seafloor [11, 12]. Many previous studies have employed benthic grabs and cores, sediment profilers, or sediment surface photography to investigate the effects of dredge spoil deposition on benthic community composition and species density [e.g. 13–15]. Laboratory experiments have also been employed to test tolerance of fauna to burial scenarios [16–18] and for short and long-term effects of suspended sediments and pollutant chemistry on survival [11, 19]. Negative impacts and community recovery rates to reference or pre-impact conditions were found to vary, with leading explanatory variables including sediment type (especially grain size and toxicity), deposition frequency and cumulative depth, and the nature of the pre-impact community (opportunistic/ephemeral versus stable/long lived). While some infaunal communities are extinguished by sediment deposition events, in others organisms are adapted to sediment disturbance and are capable of reestablishing respiratory pathways or escaping from relatively deep burial (10s of cm). In contrast, more sedentary sessile epifauna (especially suspension feeders), often fare poorly even at shallow deposition depths [16, 17]. Investigations of the direct effects of deposition events on mobile epifauna are rare because of methodological constraints, and *in situ* studies of sediment discharges on epifauna have not been previously reported.

Deposition events from hopper dredges occur in three phases [20]. The first, “convective descent”, is the release, mixing, and descent of a sediment-water plume through the water column. “Dynamic collapse” occurs as the plume encounters the seabed, where it spreads laterally and decelerates. If the dredge transits along the disposal transect, the dynamic collapse manifests as a lateral surge propagating along the seabed from the impact track. As the slurry loses momentum, the sediment is deposited and resorted by the hydrodynamic regime in a process known as “passive transport and diffusion”. These events are highly energetic. Modeling indicates velocities of the sediment-laden slurry at the seabed can reach 3-4 m/s [21], subsequently verified by near bottom vector velocimeter measurements [22]. However, the lateral deposition footprint at shallow nearshore sites may be limited to ~100 m from the ship track centerline, depending on sediment characteristics [e.g. 21, 23]. Our study of impact events measured aspects of the dynamic collapse and passive transport and diffusion phases.

To minimize some of the potential negative effects of sedimentation and the lateral surge on benthic fauna, the MCR beneficial sediment program employs a “thin-layer” dispersal technique to limit sediment mounding that could harm fauna and negatively impact wave amplification and navigation. In thin-layer disposal, sediment is gradually released while the vessel transits a predetermined disposal track [24]. Disposal tracks are arranged to distribute sediment widely within the prescribed beneficial use site. Together, these procedures aim to reduce the sediment depth of individual disposal runs and also disperse the cumulative loads over an extended area. At the MCR, individual deposition events average around 4,200 m^3^ and were expected to result in a footprint of approximately 1.8 × 10^5^ m^2^ and sediment depth range up to 9 cm along the centerline [23]. Sediment deposition exceeding this depth could cause mortalities to benthic organisms such as the Dungeness crab [25, 26]. These model results derived from scaling calculations could only be approximated in the experimental test facilities used by Johnson and Fong [20] and Vavrinec et al. [26], and thus *in situ* experiments and observations were required for confirmation.

Our experiments utilized underwater video to measure the acute effects of sediment deposition events *in situ*. “Acute effect” is defined as any disturbance (movement, displacement, burial, mortality) at impact treatments compared to control treatments. The experiments utilized baited benthic video landers (BVLs) in a before-after control-impact (BACI) design that compared the impact of the sediment plume on organisms attracted to bait to a non-impact control treatment. We had two main objectives: 1) Characterize the impact plume and sediment dynamics, and 2) Ascertain acute effects of deposition on the Dungeness crab. We also comment on observations of other organisms and their interactions.

## 2.0 Methods

### 2.1 Study site and sediment disposal

Our primary research area was the South Jetty Site (SJS) in the nearshore ocean off Clatsop Beach, Oregon USA (Fig 1), within the Clastop Plains subcell of the Columbia River Littoral Cell [27]. The 6.2 km^2^ experimental beneficial use site was comprised of relatively level, sandy substrate located between 14 and 16 m depth. The sediment deposited at SJS is clean medium-fine sand (0.13-0.35 mm diameter with < 3% fines) dredged from the nearby Columbia River navigation channel, and is of similar constitution to sediments at the disposal site (D_50_ = 0.19 mm with < 3% fines) [7]. Sediment deposition runs were conducted by the USACE ship *Essayons*, a multiple-door hopper dredge. Limited observations were also made at the Shallow Water Site (SWS) on the north side on the Columbia River channel and at the Deep Water Site (DWS), located 5 km offshore at 70-90 m depth that comprises the primary ocean dumping side for Columbia River dredge deposits [28].

**Fig 1.**
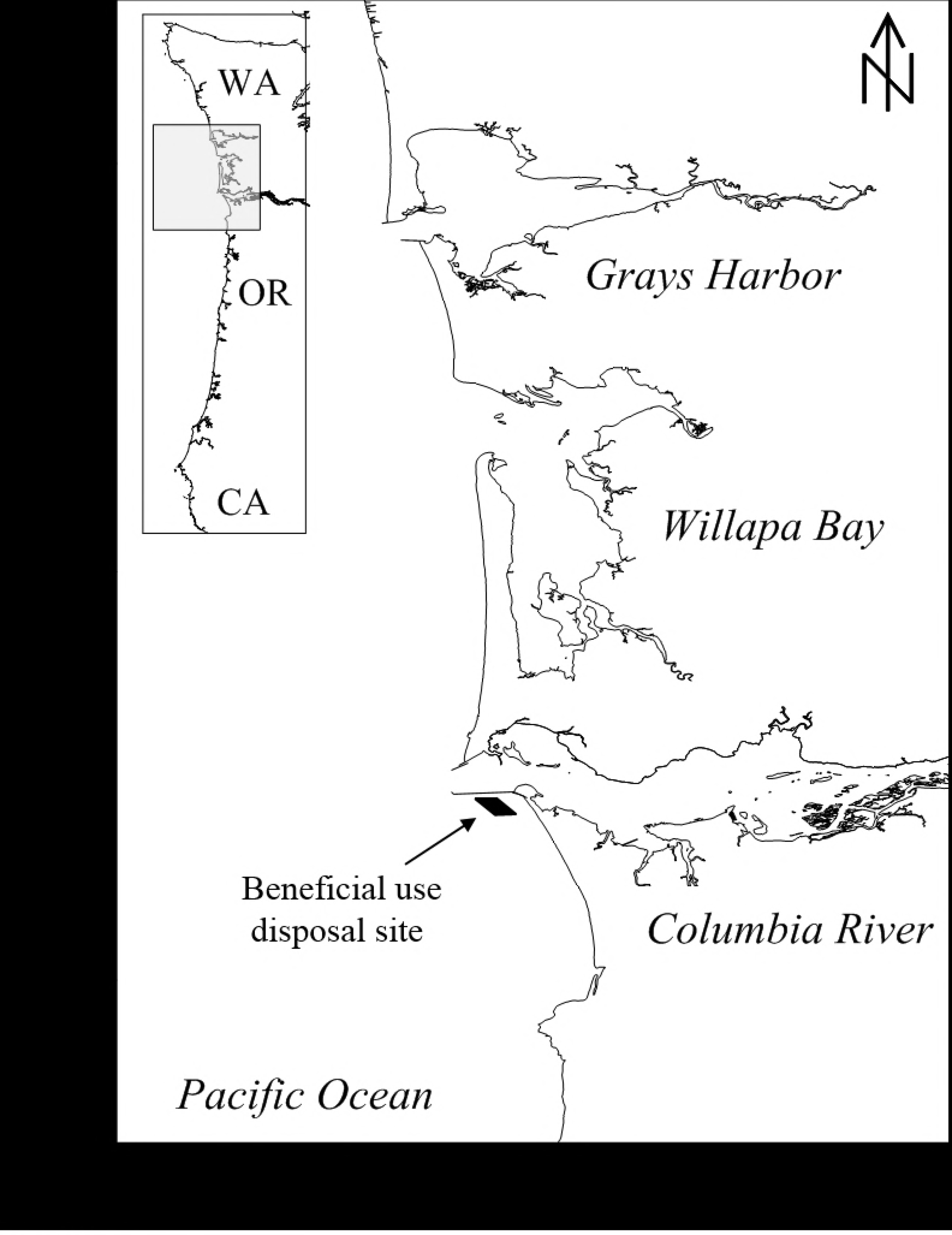
Study region at the mouth of the Columbia River. Location of the South Jetty Site beneficial use area is indicated. Inset provides a regional perspective in the Pacific Northwest of the USA.

### 2.2 Benthic video landers

We built a series of BVLs to measure acute effects of sediment deposition on epifauna. These platforms were deployed immediately before disposal events and performed two functions: First, they measured sedimentation levels, and second, they recorded imagery showing the direct effects of sediment plumes on organisms.

Each BVL had a base consisting of a 0.5 m^2^ circular rim made from 15 mm stainless steel and slatted with a series of flat metal strips that prevent burial in the substrate (Fig 2). Weights were attached to the rim of the base for stability. Welded to the base was a 110 cm central pole with four curved support ribs. The central pole held a downward looking video camera (GoPro Hero, models 2 or 3+) and an underwater light source (Intova IFL WA ZOOM). The downward viewing camera captured images of the 0.5 m^2^ base area that was used to estimate sediment cover from deposition events, and to determine time series of faunal density. The underwater light source was used to evaluate the intensity of the sediment plume. A second, inwardly looking camera imaged sediment accumulation against a 200 mm staff gauge located on the bottom of the central pole, and also aided in behavioral observations. Each BVL was baited with diced northern anchovy (*Engraulis mordax*) placed in a perforated plastic container secured to the base.

**Fig 2.**
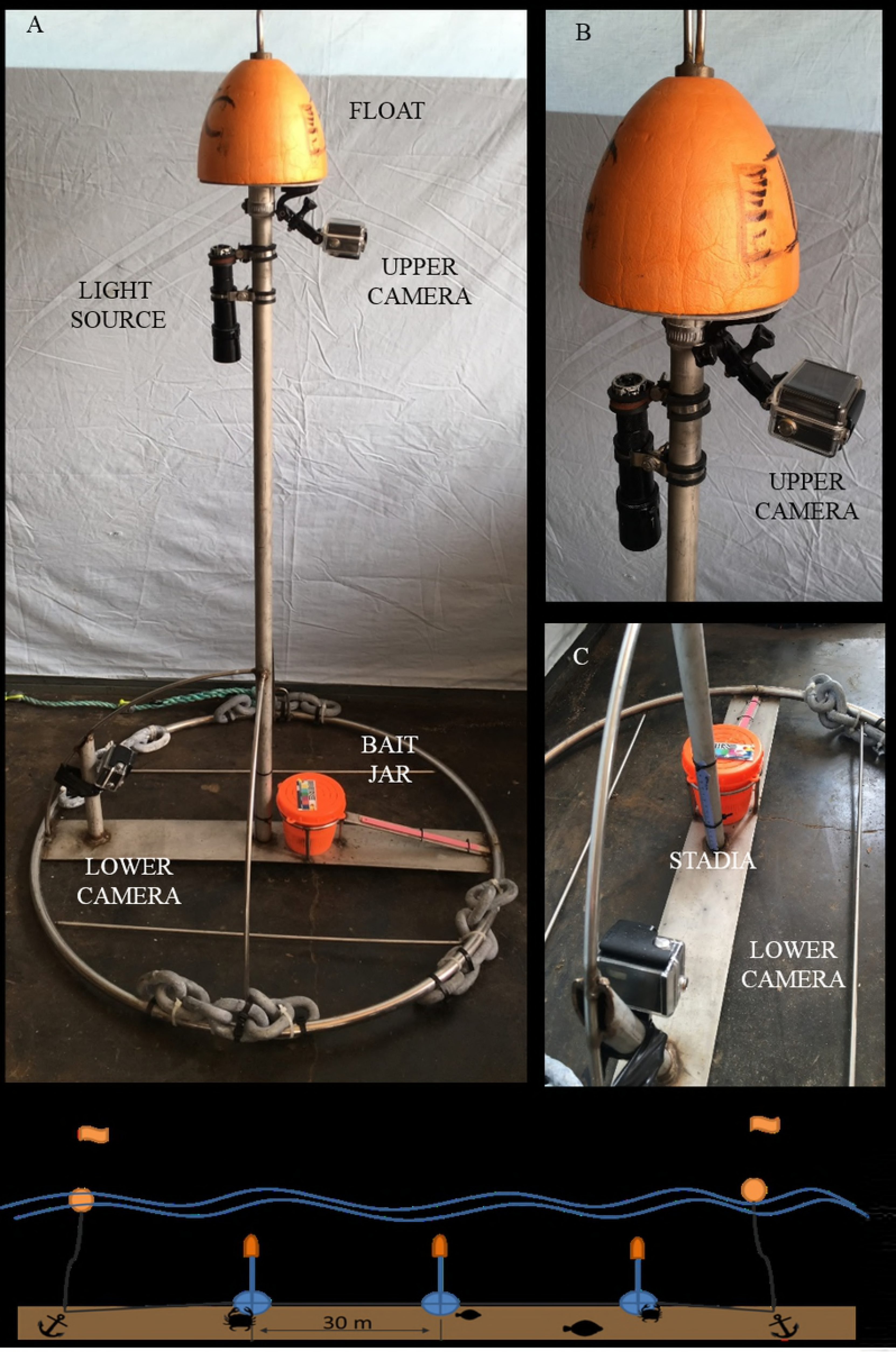
Design of benthic video lander (BVL) indicating position of upper and lower cameras, light source, sedimentation stadia, and bait jar. (A) Overview. (B) & (C) Details of upper and lower camera positions, respectively. (D) Schematic of BVL units on “daisy chain”.

Video data were recorded at a standard high definition resolution of 1920 × 1080 pixels at 30 frames/s on the “wide” setting (1080p 30 wide). This setting provided the best balance between resolution, field of view, and battery life (which was ~ 2.5 h with a standard battery at full charge). In 2016 we added extended battery packs to lengthen the post-impact observation period.

Three replicate BVLs were deployed at Control and Impact sites to test the null hypothesis of no acute effect of the lateral plume on crab density. At the Impact site, individual BVLs were joined in a “daisy chain” configuration that allowed the dredge to release sediment directly over the BVLs without fouling mooring lines. There were 30 m between BVL units, thus assuring the Impact replicates would be within the sediment plume impact zone. Individually deployed BVLs were deployed at the Control site, and during 2015 and 2016 as a continuation of the daisy chain line at the Impact site. These distal BVLs were deployed 100 m from centerline of the disposal track (70 m from the outer BVLs), and were used to gauge effects lateral from the targeted deposition track.

During an experiment, BVLs were deployed at Control and Impact locations (separated by ~1.9 km), and the dredge *Essayons* then transited a pre-determined course over the Impact BVL daisy chain. This period between BVL deployment (T_0_) and sediment impact (T_I_) was between 30 and 45 min, depending on coordination with the dredge vessel, and allowed organisms attracted to the bait to accumulate at the BVLs. No sediment impact effects occurred at the Control site. BVLs were retrieved after batteries were deemed to be exhausted after at least 2.5 h. Note that crab acoustic tagging experiments were often initiated during the BVL deployments [29].

### 2.3 Video analysis

We used the timing of the sediment plume at each impact BVL to define pre- and post-impact time periods. Pre-impact sequences ended once the sediment plume crossed the perimeter of the BVL base. Post-impact sequences began consecutively, ending either when the BVL was retrieved or more often when the camera battery was exhausted. Between coordinating deployment of the cameras with the dredge vessel and variable battery life of the camera batteries, post-impact video sequences ranged from 9 to 75 min in 2014 and 2015 and up to 120 min in 2016 with extended battery packs. All video footage was sequenced and edited in Adobe Premiere Pro CC or CyberLink PowerDirector 13. See Fields [28] for further details.

#### 2.3.1 Observation of sediment levels

To assess potential for organism burial, we examined the upper and side video imagery from the Impact treatment immediately following dissipation of the sediment plume. With the upper camera we estimated the percent of the 0.5 m^2^ base that was covered with sediment. With the side camera we measured the depth of deposited sediment from the staff gauge. The results are tabulated.

#### 2.3.2 Organism density

Each BVL video was scored at five minute intervals for organism density (D_T_). We concentrated on Dungeness crab (*Cancer magister*) and dog whelk (*Nucella spp*) because they were the most abundant invertebrate organisms and exhibit contrasting locomotor abilities. Species were enumerated only when observed contacting or within the 0.5 m^2^ base of the BVL. Note that counts were not possible during impact events, when visibility was reduced to zero. These periods proved to be less than seven minutes in duration and were typically approximately three minutes in duration.

To test for impact effects, we standardized the data in two ways. First, because maximum organism densities (D_MAX_) between BVLs could vary (for crab, 2 to 20 ind / m^2^) between and within experiments, we normalized densities as proportion of D_MAX_ (D_N_ = D_T_ / D_MAX_). These data comprised the independent variable for statistical tests. Second, the time from deployment to the sediment deposition event (T_I_) varied from 20 to 70 min among experiments (Table 1), and Control and Impact treatments were deployed up to 20 min apart. To standardize the time of organism presence at Control and Impact treatments, we calculated the mean T_I_ among Impact replicates (which were within 5 min of each other), and applied that value to the Control time series, thus standardizing the crab accumulation period within an experiment. This process resulted in four treatment groups (dependent factors) for each experiment: Control-Pre, Control-Post, Impact-Pre, and Impact-Post. Within an experiment, the total length of the “Pre” treatments were equal, but the “Post” video lengths were different due to variation in battery usage.

**Table 1.**
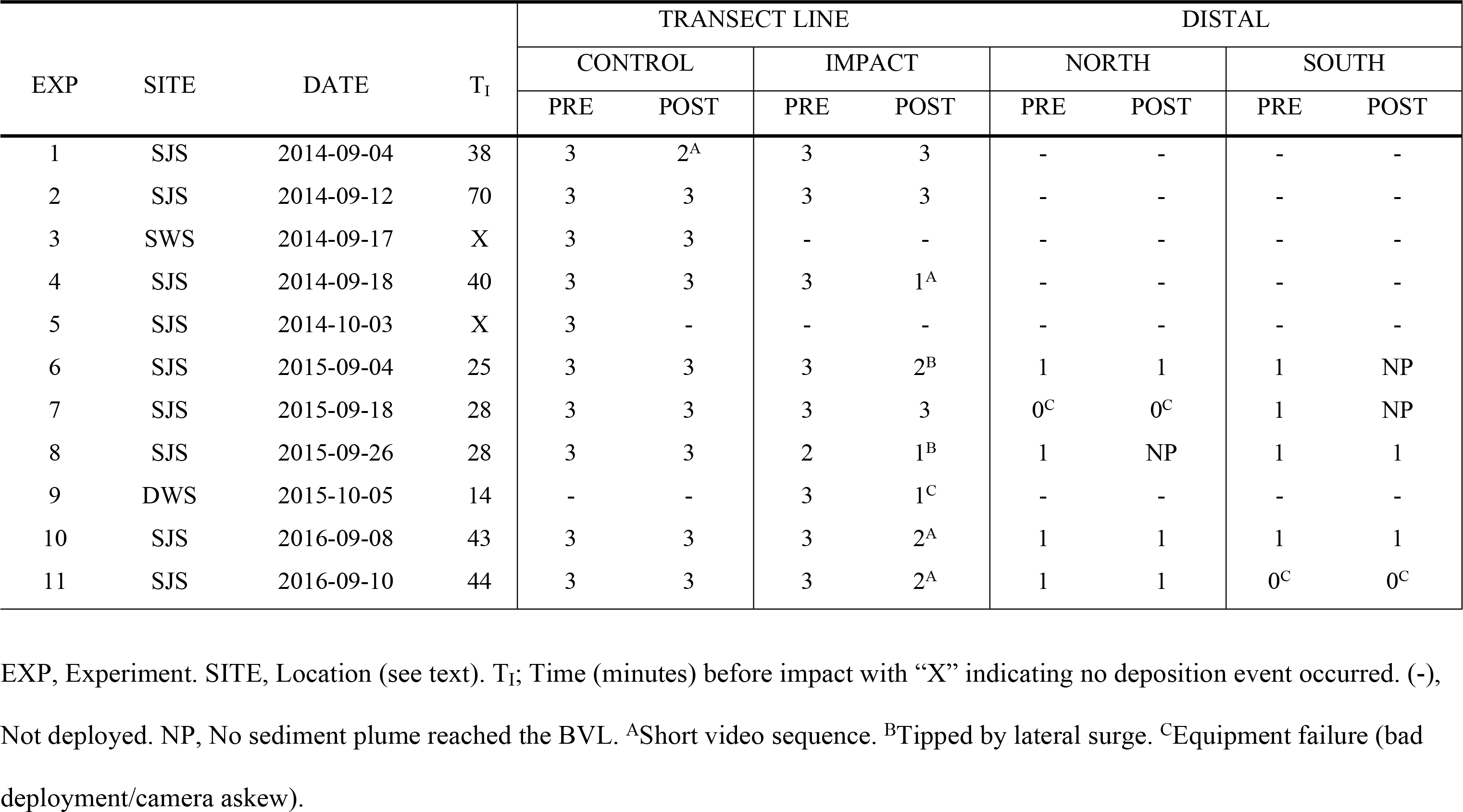
Summary of BVL experiments indicating number and status of replicates for each treatment group. There were three replicates along the Transect Line but only one for Distal BVLs.

To test for acute impact effects on mobile epifauna, we calculated the mean D_N_ in three time steps before and three time steps after T_I_ for each BVL time series (15 min each before and after T_I_). We then ran a two-factor Analysis of Variance testing the significance of D_N_ by Experiment × Treatment Group. Tests were executed separately for crab and whelk.

To further explore the potential impacts of the sediment plume, we determined the time of first return (T_R_) of crab during the post-impact period of the impact BVLs. We compared BVLs directly under the dredge track to those deployed distal to the main deposition track. Regression was used to examine the reliance of return time to length of video sequence; otherwise results are descriptive because there were no replicates in the distal treatments.

## 3.0 Results

During 2014, we deployed BVLs three times at SJS during sediment deposition events (Exp 1, 2, and 4) and one time during the recovery period when no deposition events occurred (Exp 5). We also made deployments at the Shallow Water Site for observation purposes when no direct impact event occurred (Exp 3). During 2015 we deployed BVLs at SJS an additional three times (Exp 6-8), and tested the BVL technology at the Deep Water Site (DWS) in 70 m of water during a deposition event (Exp 9; data not shown). In 2016 we made two final experiments at SJS with extended battery packs to further investigate return of crabs after dredged sediment impact (Exp 10-11). Table 1 denotes details on the experiments and the number of usable replicates.

### 3.1 Sediment deposition dynamics

Along the dredge transect line, videos revealed deposition events resulted in a rapidly moving sediment plume that enveloped the BVLs (Figs 3 and 4; Video 1.1 and 1.2). This lateral surge had horizontal current velocities of 2.4-3.0 m/s as estimated from particle trajectories [23], and video data blacked out during deposition events, an indication of high levels of suspended sediment. However, the impact period was relatively brief. Based on clarity of the image from the downward-looking camera, the impact of the sediment plume did not exceed seven minutes.

**Fig 3.**
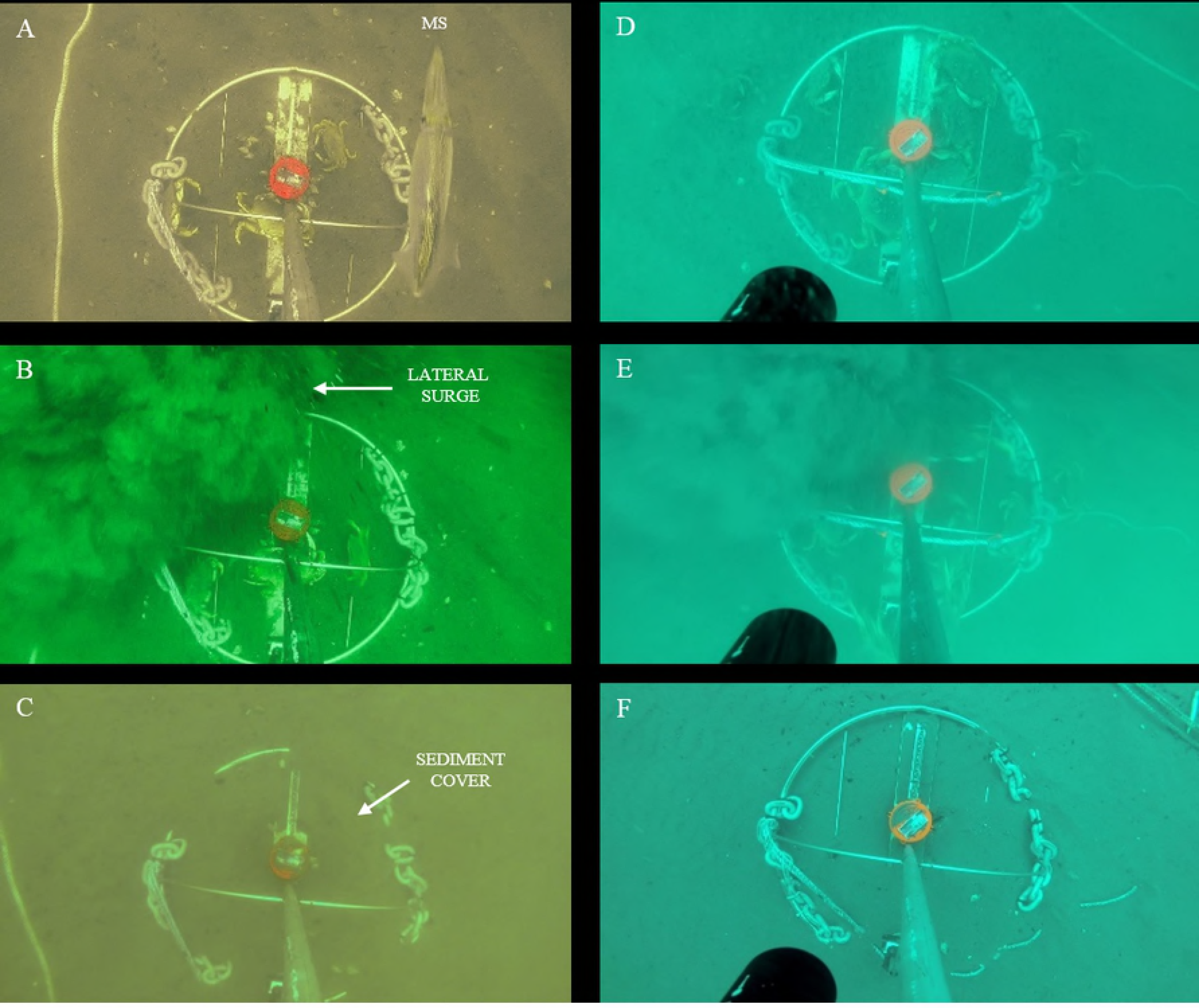
Plan view examples of frame grab images illustrating before, during, and after impact of the lateral surge of a disposal event at BVLs. Images were 2 to 5 min apart in the video sequence. (A)-(C) Deployment on 2016-09-08. (D)-(F) Deployment on 2016-09-10. MS; the market squid *Doryteuthis opalescens*.

**Fig 4.**
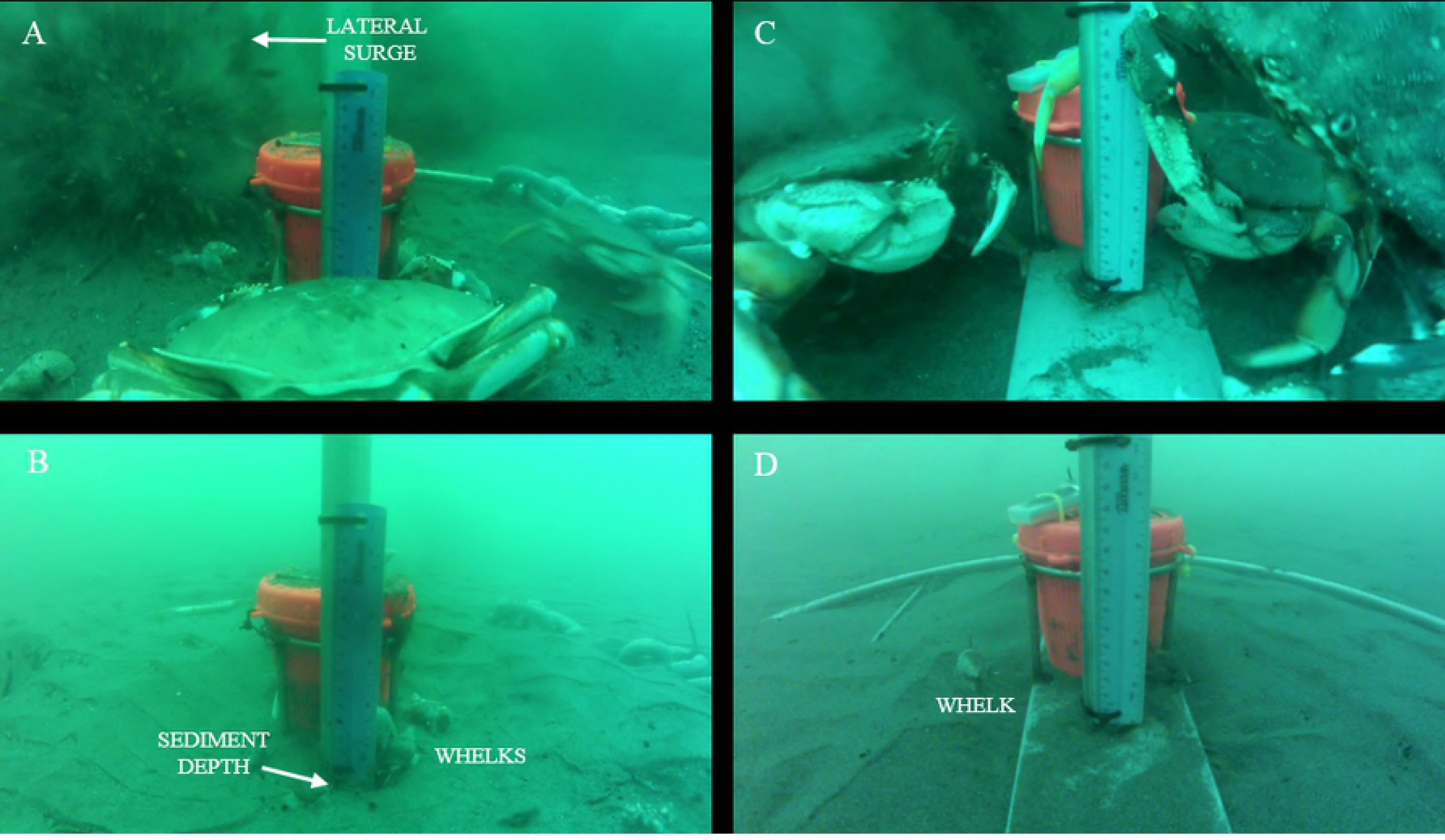
Profile view examples of frame grab images illustrating during and after impact of the lateral surge at BVLs, indicating contrasting sedimentation levels. Note presence of whelks remaining on the newly deposited sediment surface while Dungeness crabs are dispersed. (A) & (B) Deployment on 2016-08. (C) & (D) Deployment on 2016-09-10. V01.1 – V01.2 Examples of the lateral surge and subsequent sedimentation levels at BVLs during Exp 02 (2014-09-12) and Exp 10 (2016-09-10). Plan and profile video sequences are presented, with all imagery processed at 5x actual speed. BVL-N, -C, and -S refer to North, Center, or South position of each benthic video lander on the daisy chain mooring.

Mean percent sediment cover deposited on the BVLs along the dredge transect line was 86.9% ± (range 45 to 100%) (Table 2). Overall deposition levels averaged 1.1 cm ± 1.0; range <0.5 to 4 cm), and levels often did not register on the measurement scales. At the distal BVLs, ~ 100 m from the transect centerline, mean percent cover was only 22.5% ± 25 SD (range 0 to 60%;), and on three occasions the impact plume failed to reach one of the paired BVLs. These observed deposition levels would not result in significant organism burial.

**Table 2.** Sedimentation on Impact BVLs. Sediment cover is percent of 0.5 m^2^ base with visible deposits measured with top camera. Sediment depth is cm of sediment deposited on measuring stadia determined from the side camera. Side cameras were not deployed on distal BVLs.

| EXP | DATE | SEDIMENT COVER (%) |  |  |  |  | SEDIMENT DEPTH (cm) |  |  |
| --- | --- | --- | --- | --- | --- | --- | --- | --- | --- |
|  |  | TRANSECT LINE |  |  | DISTAL |  | TRANSECT LINE |  |  |
|  |  | NORTH | CENTER | SOUTH | NORTHF | SOUTHF | NORTH | CENTER | SOUTH |
| 1 | 2014-09-04 | 90 | 100 | 100 | - | - | <0.5 | <0.5 | <0.5 |
| 2 | 2014-09-12 | 90 | 75 | X | - | - | 0.8 | <0.5 | X |
| 4 | 2014-09-18 | X | X | 50 | - | - | X | X | <0.5 |
| 6 | 2015-09-04 | 60 | X | 100 | 10 | NP | 1.5 | X | 1 |
| 7 | 2015-09-18 | 100 | 90 | 90 | 50 | NP | 4 | <0.5 | 1 |
| 8 | 2015-09-26 | 45 | X | X | NP | 10 | 1 | X | X |
| 10 | 2016-09-08 | 100 | 100 | 100 | 60 | X | 1 | 3 | 1 |
| 11 | 2016-09-10 | 75 | 100 | 100 | 10 | X | <0.5 | X | 1 |

(-); no Distal treatment deployed. NP; no plume observed. X; deployment failed.

### 3.2 Organism density during sediment plume impacts

The primary organisms quantified from BVL deployments were Dungeness crab, dog whelk (*Nucella* spp), hermit crab (Paguridae) and several species of benthic fishes (full species list in Fields [28]). However, only Dungeness crab and whelks were abundant enough for further analysis. This was partially due to antagonistic behavior by crabs toward other organisms, excluding them from the BVL base area. Requirements for the BACI design were fulfilled for six experiments with Dungeness crab. For whelk, the BACI design was fulfilled for only one experiment, and we therefore conducted single factor ANOVAs for four experiments which had sufficient data (as detailed below).

For the Dungeness crab, results were consistent between experiments, and were significantly different among treatment groups (Fig 5). Mean proportional density of crab was similar between Control and Impact locations during the pre-impact period (Fig 6), and densities tended to increase to an asymptote 20 to 30 min into the deployment. Fluctuations in the mean number occurred because crabs actively moved in and outside of the BVL base rather than accumulate as they would in a standard crab pot. When the lateral surge swept over the Impact site, all crabs were displaced (Figs 3 and 4; Video 1.1 and 1.2). Displacement occurred both by escape behavior, where crabs attempted to avoid the approaching sediment plume, and by entrainment, where crabs were engulfed and swept away by the plume. No crab remained at the BVLs post-impact, nor were crab buried *in situ*. Concurrently, mean normalized densities remained comparatively high at the Control site. Thus, we detected a significant effect of the lateral surge on Dungeness crab (P<0.001) (Fig 6; Table 3)

**Fig 5.**
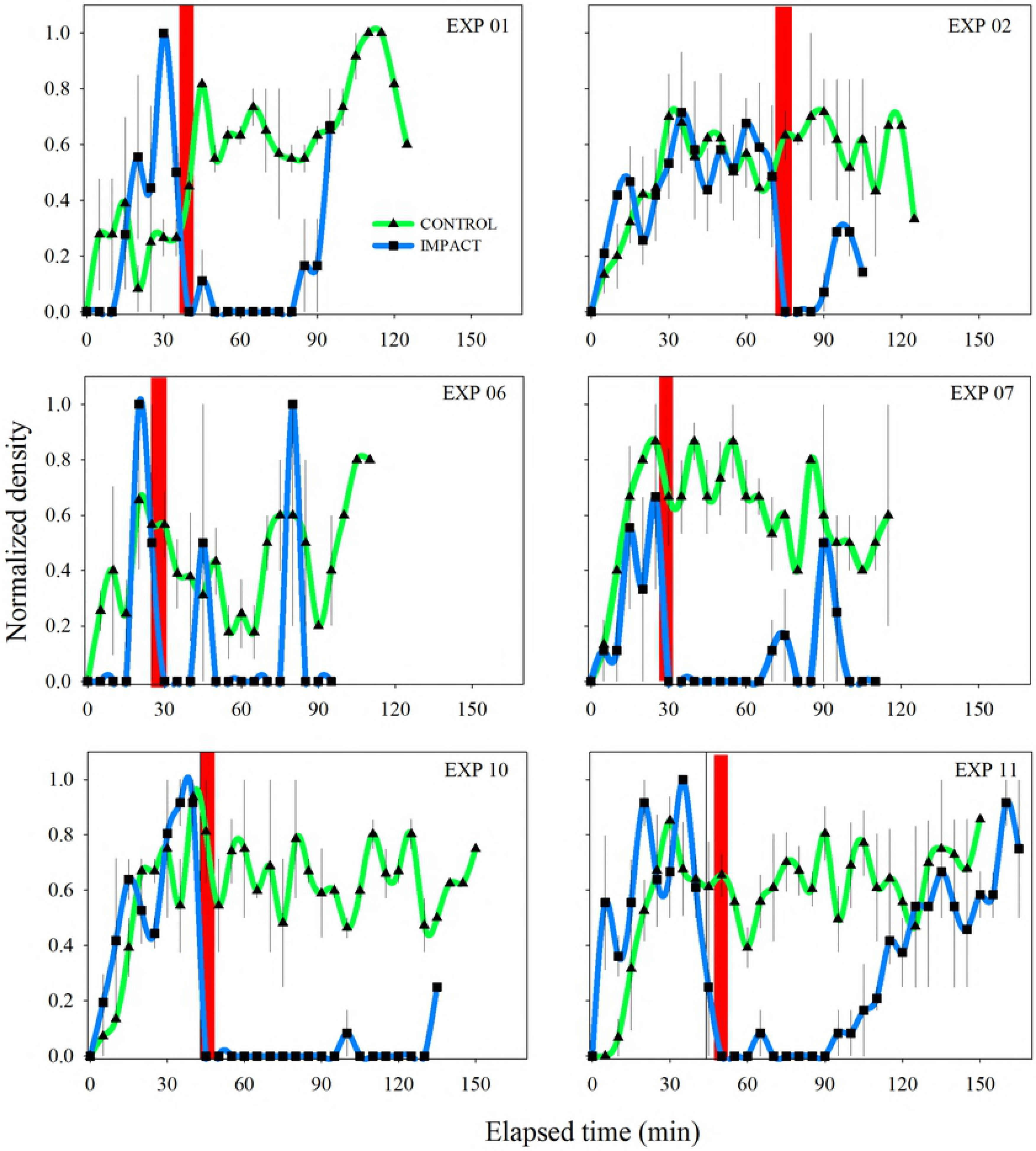
Time series of mean (± standard error) normalized density of Dungeness crab at Control and Impact sites. Red bar denotes timing of lateral surge and designates pre- and post-impact periods.

**Fig 6.**
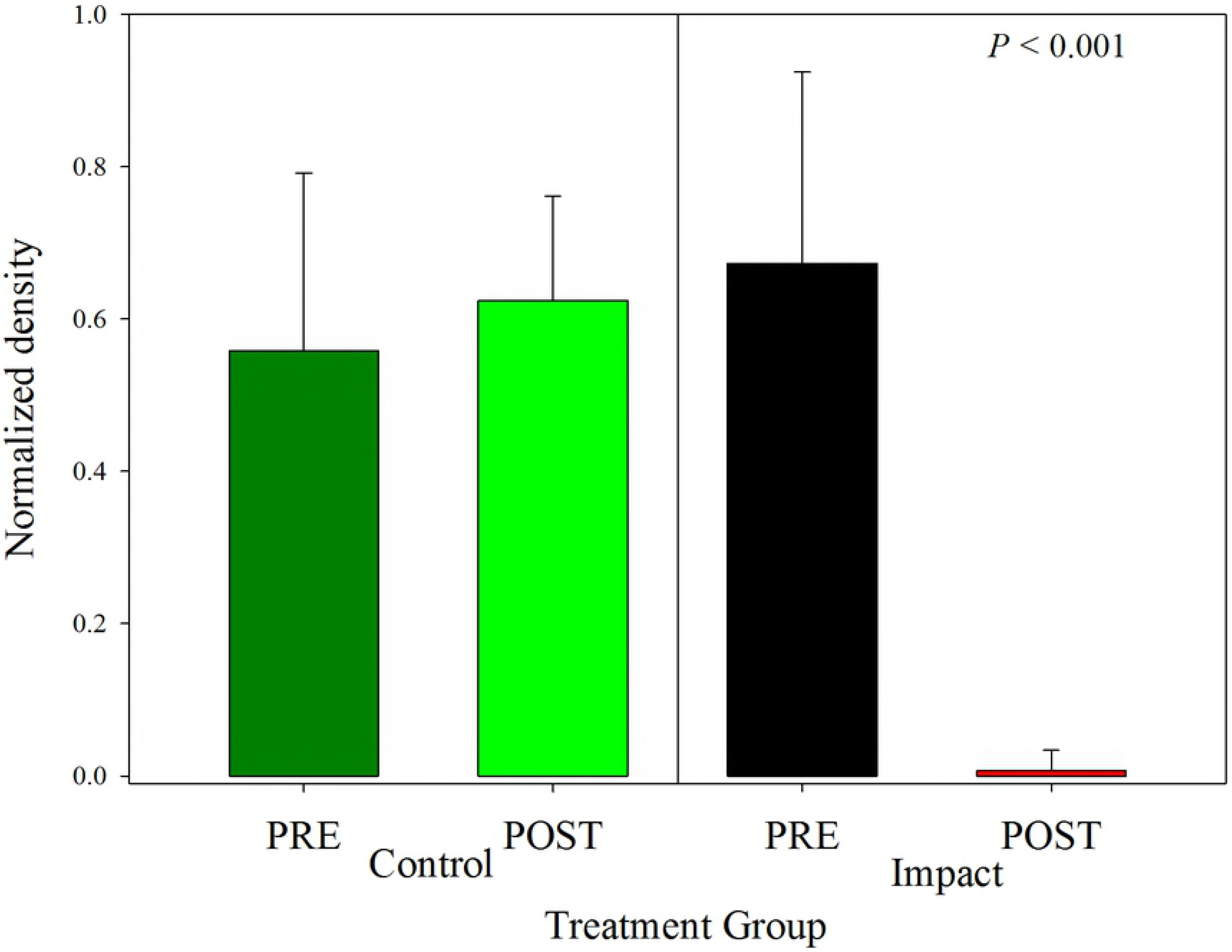
Comparison of treatment effects for mean (± standard error) normalized density of Dungeness crab. Densities were determined 15 min before (pre-) and after (post-) arrival of the lateral surge at Control (C) and Impact (I) sites. The post-Impact treatment was significantly different from the others.

**Table 3.** Two-factor Analysis of Variance testing Dungeness crab normalized densities (D_N_) among pre-and post-deposition at control and impact sites

| Effect | SS | DF | MS | F | p |
| --- | --- | --- | --- | --- | --- |
| Intercept | 13.686 | 1 | 13.686 | 454.416 | <0.001 |
| EXP | 0.325 | 5 | 0.065 | 2.157 | 0.077 |
| TRT | 4.675 | 3 | 1.558 | 51.743 | <0.001 |
| EXP TRT | 0.537 | 15 | 0.036 | 1.189 | 0.317 |
| Error | 1.265 | 42 | 0.030 |  |  |

SS; Sum of Squares. DF; Degree of freedom. MS; Mean Square. F; F-statistic. P; Probability ( $\alpha=0.05$ ). EXP Experiment (see Table x). TRT, Treatment (Control-Pre, Control-Post, Impact-Pre, and Impact Post). Tukey post hoc indicates Post-Impact < other treatments, which are not different.

In the post-impact period, crabs often were observed to return to the sediment covered BVL (Fig 7). The mean time for a crab to reoccupy a BVL was relatively variable (23.2 min ± 28 SD) and was affected by the length of the post-impact video length. Of 16 trials with BVLs on the transect line, six had no returns up to 90 min post-impact. For the remainder, time of return was moderately related to length of the video record (r = 0.40), suggesting increased occupancy over time. The distal BVLs generally experienced a diminished impact plume, and all four of the trials that the lateral surge impacted had returns within 10 min. These data put boundaries on the detrimental effects of sediment deposition events and suggests a limited acute effect on crab foraging patterns.

**Fig 7.**
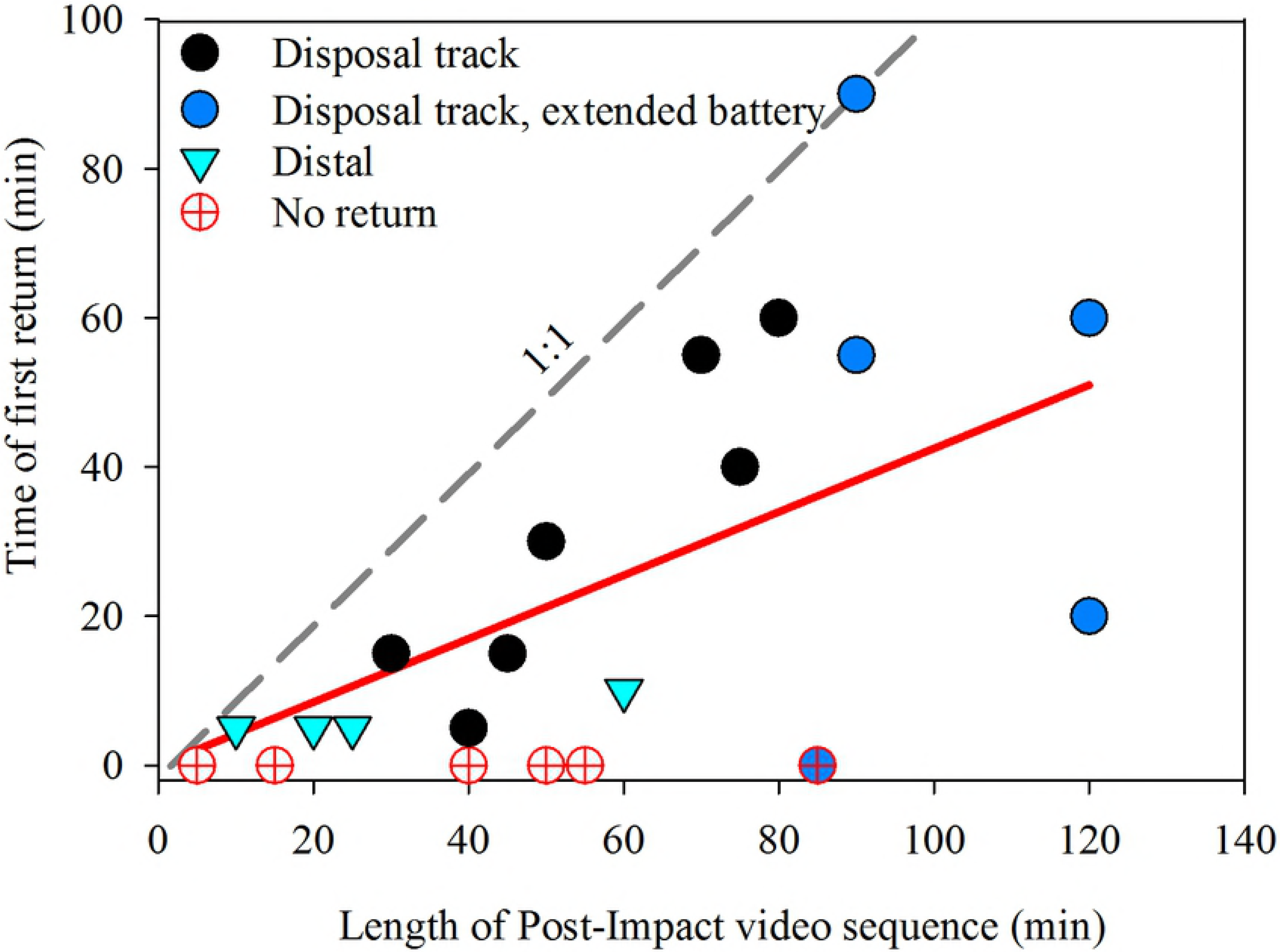
Time of first return of Dungeness crab to BVLs following a sedimentation event by video length (= elapsed time). If “no return” values are neglected, correlation increases to *r* = 0.49.

Evaluation of impact effects on whelk was hampered by patchy and inconsistent densities between Impact and Control treatments during 2015 and by failure of whelk to be observed at impact sites at pre-impact treatments in both 2014 and 2015 (Fig 8) Whelk were not quantified in 2016. These conditions violated the BACI design. We therefore ran individual one-factor ANOVAs testing variation between Pre- and Post-Impact proportions without a Control comparison. Of the four statistical tests conducted, three were non-significant and the final test indicated higher densities in the post-impact treatment (Fig 9, Table 4). Inspection of the time series in Fig 8 indicates overall densities were not consequentially affected by the lateral plume and that whelk densities tended to remain high at the BVL during the Post-Impact period.

**Fig 8.**
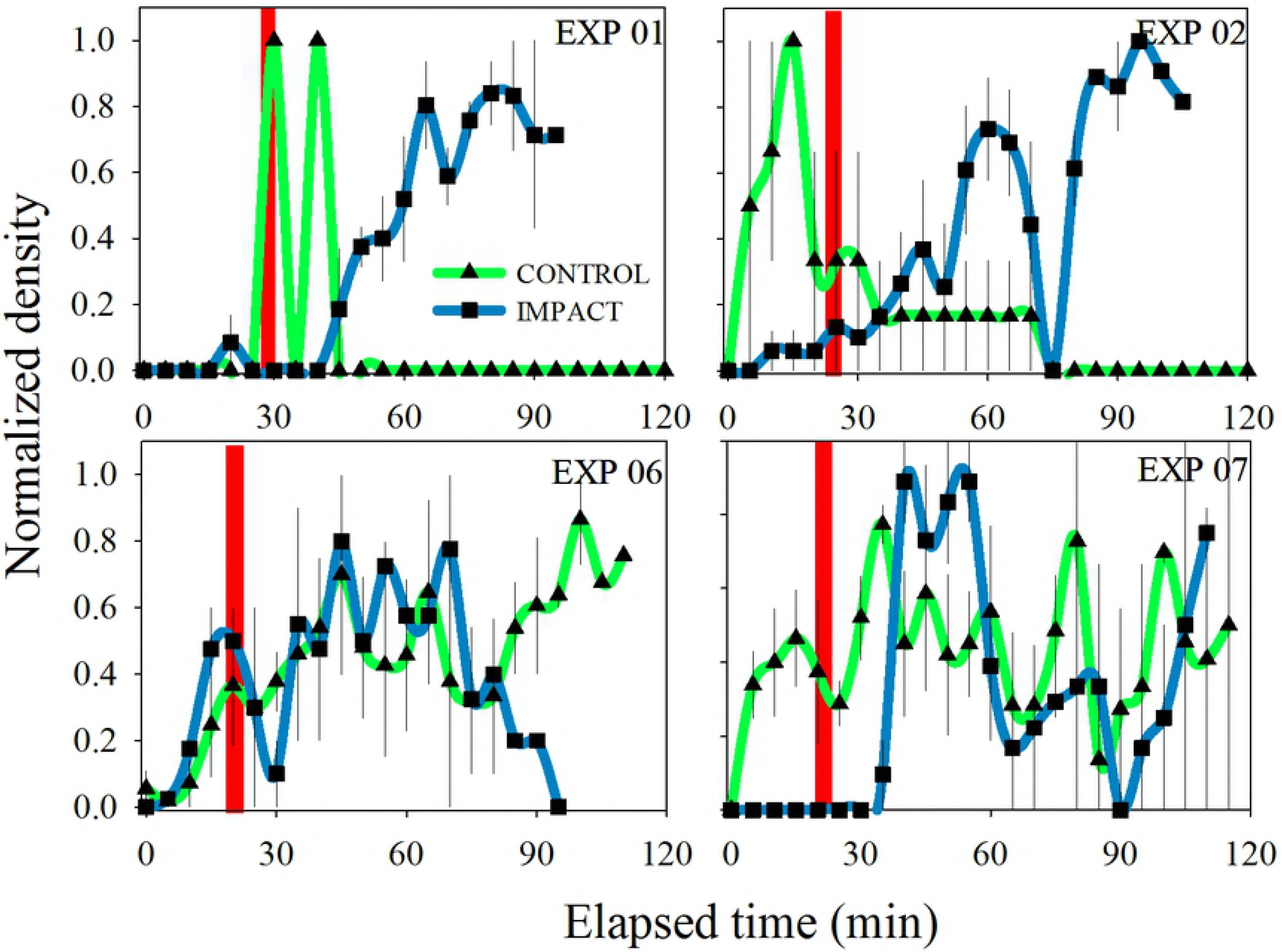
Time series of mean (± standard error) normalized density of whelk at Impact and Control sites. Red bar denotes timing of lateral surge.

**Fig 9.**
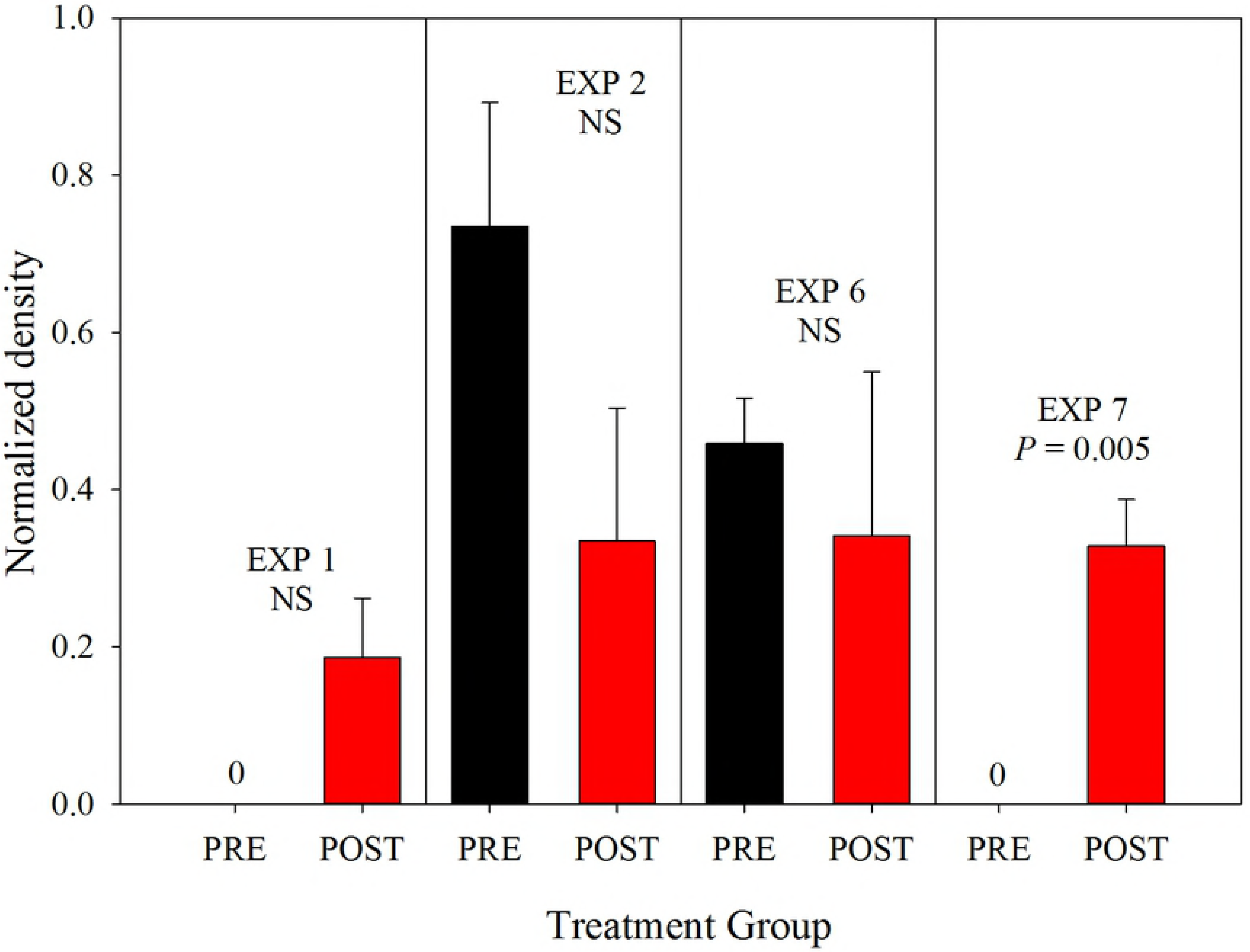
Comparison of treatment effects for mean (± standard error) normalized density of whelk. Treatments were before (pre-) and after (post-) arrival of the lateral surge at the Impact site. Oneway ANOVAs were used to test for significant differences.

**Table 4.** One-factor Analysis of Variance testing dog whelk normalized densities between pre- and post-deposition periods at impact sites.

| EXP | SS | DF | MS | F | P |
| --- | --- | --- | --- | --- | --- |
| 1 | 0.034 | 4 | 0.009 | 6.119 | 0.069 |
| 2 | 0.321 | 4 | 0.080 | 2.996 | 0.159 |
| 6 | 0.094 | 2 | 0.047 | 0.291 | 0.644 |
| 7 | 0.021 | 4 | 0.005 | 30.268 | 0.005* |

SS; Sum of Squares. DF; Degree of freedom. MS; Mean Square. F; F-statistic. P; Probability ( $\alpha=0.05$ ). \*I-Post > I-Pre.

### 3.3 Qualitative observations of BVL deployments at the SWS

We conducted a BVL deployment (Exp 3) at the Shallow Water Site in 2014, a high energy location on the Columbia River bar. This experiment compared densities of crab and whelk at BVLs during a high wave period compared to more sheltered conditions at the SJS (Exp 5). Neither of these deployments included impacts from a sediment deposition event, and thus serve as observations rather than experiments. Fig 10 illustrates that occupation of crabs at BVLs was similar at all three sites, increasing within 20-40 min to an asymptote. Whelk had similar abundance patterns at SJS but were uncommon at SWS (data not shown). However, video observations of crab at SWS reveal crab agility in oscillatory flow (Video 2). The surge from waves at the SWS resulted in 50-70 cm/s bottom currents accompanied by resuspension and transport of sand. Crabs were observed to resist dislocation by remaining near the sediment surface, but those dislodged by the more energetic bursts of water and sediment often returned to the BVL within a few seconds. Foraging crabs were also observed to ascend into the water column and travel several meters before dropping back to the substrate, and repeated sojourns could result in directional movement toward the BVL. Crabs did not bury to avoid near-bottom currents. These observations of motility in naturally energetic conditions contrast behaviors observed during the unidirectional force exerted by deposition events, as discussed below.

**Fig 10.**
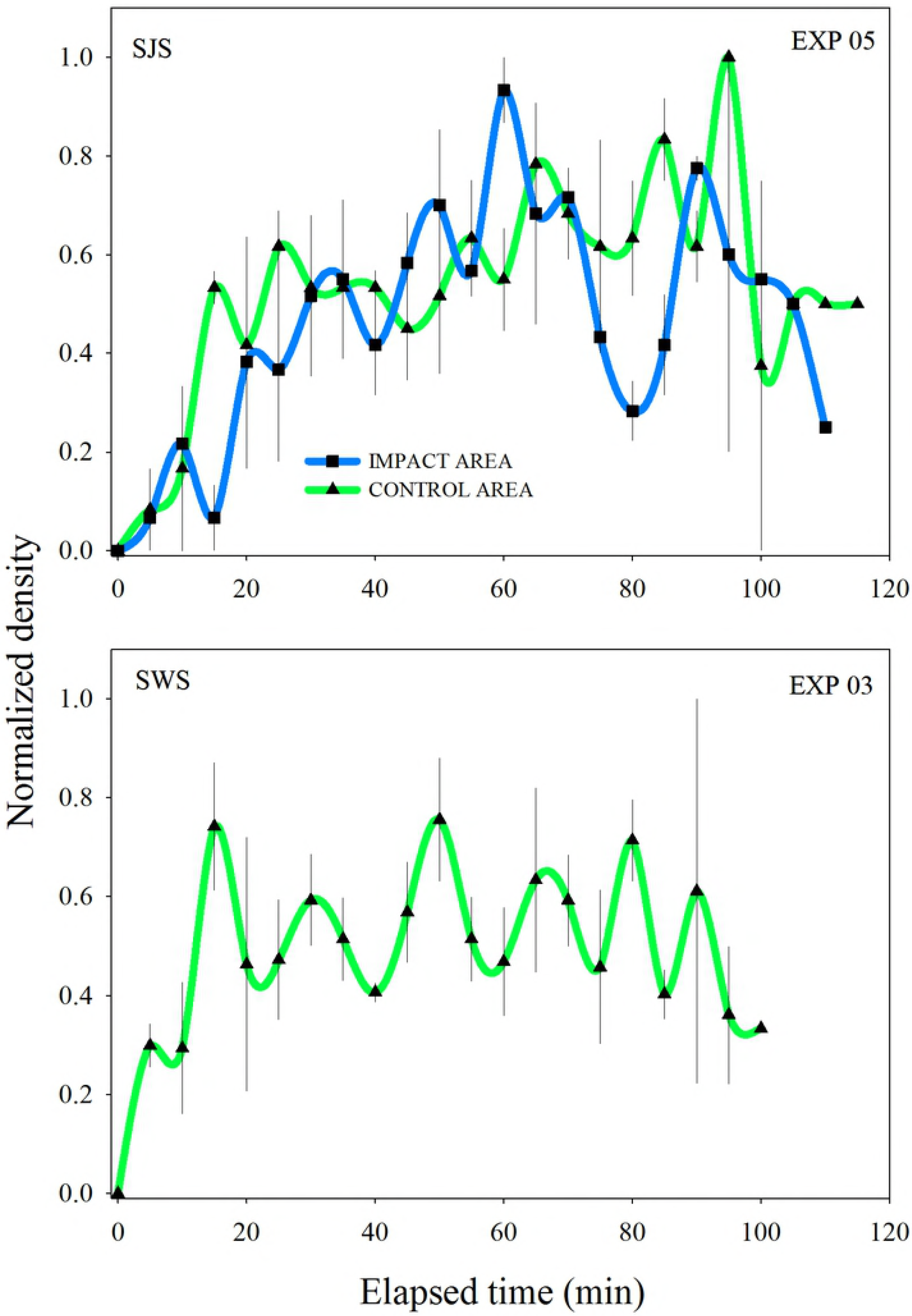
Time series of mean (± standard error) normalized density at Impact and Control areas. (A) Dungeness crab at the South Jetty Site on 2014-09-17. (B) Dungeness crab on the Columbia River bar at the Shallow Water Site on 2014-10-03. The times series at (B) occurred in heavy swell. Neither of these deployments included impacts from sediment deposition events.

V02. Example of Dungeness crab movements in a high surge environment at the Shallow Water Site (SWS) during Exp 04, 2014-09-10.

Crabs both resisted high surge events and also navigated to the bait source within the oscillatory flow.

## 4.0 Discussion

Sediment deposition events had a significant effect on crab density at BVLs. Both our video observations and instrument measurements of near bed velocity and turbidity measurements [22] yield a complimentary assessment of the dynamic collapse and the passive transport and diffusion phases of sediment deposition. Deposition events were characterized by a sudden increase in velocity (up to 3.0 m/s from ambient velocities < 0.20 m/s) coinciding with a large increase in turbidity measured by optical backscatterance [22]. The corroborating video sequences show the approach of a turbidity front-like lateral surge and an immediate obscuring of video imagery by suspended sediments (Figs 3 and 4; Video 01). These observations characterize the dynamic collapse as an energetic front laden with sediment. However, equally striking was the rapid passing of the front during the passive transport and diffusion phase: velocity at the instrument mooring returned to ambient levels within 2 min and OBS values within ~ 5 min [22]. Similarly, high currents and turbidity levels diminished in our video imagery within 5 to 7 min. These results fit within predicted impacts based on model parameters of sediment grain size, water depth, and release rate presented by Pearson et al. [21].

Moritz et al. [23] calculated the bulk sediment concentration (sediment plus seawater) of the descending plume at contact with the seabed to be 1.3 × 10^3^ kg/m^3^, which at typical seawater density yields a suspended sediment concentration of 2.8 × 10^2^ kg/m^3^. This is a dense concentration. In comparison, Peddicord and McFarland [30], in laboratory suspended sediment challenge experiments, found a concentration of only 9.2 kg/m^3^ induced an incipient mortality to 30-40 mm Dungeness crab. However, these challenge tests used fine grained sediments (mud derived from an industrialized harbor thus possibly toxic) and had a duration of 8 d, and so were substantially different from short term impacts from sand depositions we observed. A metaanalysis conducted by Wilber and Clark [11] concluded few detrimental effects on crustaceans would be expected at suspended sediment concentrations and relatively short time intervals caused by dredging operations. In agreement, our results and that of Golder [22] indicate the high settling velocity of fine sand sediments at the MCR sites and the overall dispersive nature of the wave climate at these nearshore sites leads to ephemeral turbidity effects from dredge deposits.

The force of the lateral surge is another matter to consider. The instantaneous bulk mass flux (bulk concentration × velocity) of the slurry impacting crabs can be estimated to be between 2.6 to 3.9 × 10^3^ kg/m^2^/s. These bulk fluxes, driven mostly by water movement, rival those estimated from submarine turbidity currents, albeit at a much smaller spatial extent (e.g [31–34]). In comparison, sediment fluxes from breaking waves in nearshore environments are generally less than 5 kg/m^2^/s [35, 36], although fluxes may be considerably higher in large storms. Thus one should expect a disruptive effect of these sediment disposal events on epifaunal organism status and behavior. However, the Dungeness crab was found resilient to simulated dredge activity conditions by Vavrinec et al. [26] who used flume experiments to examine crab responses to a discharge of sediment-laden water, and concluded behavioral responses by crab could minimize burial risk at velocities up to 3.2 m/s. Our studies of acoustically tagged Dungeness crab likewise indicate high survival and mobility after sediment deposition events [29].

We ascertained the potential for organism burial by measuring sediment deposited on the BVLs immediately after passage of the lateral surge. Post impact video from both upper and lower cameras revealed sediment deposition levels were moderate (maximum of 4 cm). Previous laboratory experiments examined the survival of crabs as functions of burial depth and time, the primary concerns being the maximum depth a crab could reestablish respiratory currents to the overlaying water or from which they could extricate themselves [25, 26, 37]. Results indicate burial up to about 10 cm was not detrimental for most crab sizes tested if they were unconstrained by experimental conditions. The sediment accumulation levels at BVLs thus were below depths of concern documented by Chang and Levings [25] and Vavrinec et al. [26].

There was a varied response by organisms to impact events. Crabs attracted to bait at BVLs often sensed the rapidly approaching sediment plume and enacted an escape response involving abrupt “scuttling” into the water column. In most cases, the plume was observed to overwhelm the fleeing crabs, and in all cases crabs were displaced from the BVLs. Gastropods and hermit crabs were impacted before they could meaningfully react, but were often not displaced from the BVL. Whelks are surface and shallow burrowing gastropods and would have little difficulty emerging from the shallow sediment layers when buried. Differences in whelk density between control and impact arrays are not yet understood, but may be related to enhanced organic deposition in the impact zone from the destruction of benthos during dredging (i.e., a positive attractant) [38]. Fish and squid in and around the BVLs generally swam from the field of view without observations of them being engulfed, so we are unsure of their fate. These behavioral observations from the video recordings are insightful for elucidating impacts on epifauna.

We observed that crabs returned to BVLs after a mean lag of about 20 min following an impact event. Other than the gastropods and hermit crabs that withstood the lateral surge, Dungeness crabs were generally the first organisms to reoccupy sites. Battery power of the cameras was not always sufficiently long to observe crabs returning, but these data suggest crab continued foraging relatively soon after a single deposition event. Together, field observations of crab behavior, force of the lateral surge, and sediment deposition levels agree with laboratory and modeling results and lend support to the hypothesis there are acute effects of disposal events on epifaunal crab distributions, but that recovery is rapid.

Conversely, juvenile, recently molted, and gravid female crabs may have increased susceptibility to the effects of burial and forces associated with the lateral surge. Crab size and reproductive state should be considered to ameliorate negative effects of deposition events on sensitive stages [3]. Based on size-frequency measurements from crab pot deployments, most crabs observed at BVLs were adults 100 – 170 mm carapace width, and densities of smaller juvenile crabs were generally lower in nearshore samples compared to estuary habitat during the period of deposition operations [39, 40] However, new recruits < 30 mm were observed during 2016 in both video and crab pot samples. Recruitment from the plankton in the MCR region occurs from April – November peaking in May-June [41] and growth to the adult stage (~ 100 mm) generally takes three years. The timing of deposition events in autumn thus could impact crabs recently recruited to the nearshore zone. Crabs of all sizes are expected to be vulnerable when molting, which reportedly occurs primarily during spring for females and from July-August for males in the northern California Current system [42, 43]. These time periods are mostly outside the late August – September time window permitted for disposal at the beneficial use nearshore ocean sites. However, in September 2016 we observed a pair of crabs in conjugal embrace at a BVL moments before an impact event, so at least some disruption of mating and injury to thin-shelled female crab may occur at the SJS. Less is known of the distribution of ovigerous females during the late fall brooding period. For coastal populations, females migrate from estuaries to brood extruded eggs in high salinity nearshore zones [44], but there is little survey data to identify habitat preferences of female crab in the ocean during this period. In an embayment of Vancouver Island, British Columbia, acoustic telemetry indicated brooding females converged and remained localized within sandy substrate at ~40 m depth before migrating to shallower water to release larvae [43]. Protection of such concentrations of brooding females should be of paramount concern, and more data is needed to identify and conserve these habitats. Also important is the recognition that the timing of these life-history characteristics (larval settlement, molting, and mating) vary latitudinally across the Dungeness crab range at present and may be in flux due to recent ocean temperature anomalies observed in the PNW [45].

Our use of BVLs in an experimental design was a novel approach to understand the effects of sedimentation events from dredge deposition events on mobile epifauna. The video imagery provided observations of the time series of forces directed on target organisms, and density data was extracted from the video sequences in a straightforward manner for use in hypothesis testing. Likewise, sediment deposition levels were determined immediately following the restoration of visibility and before hydrodynamic resorting could occur. The behavioral observations, while not quantified in the present project, offered unique insights for species interactions that affected density data. See Video 3 for an example of the effect of video speed on behavioral observations. Further, the methods are reproducible at other sediment deposition sites of reasonable water clarity, and it would be interesting to compare our results to sites where finer grained sediments that pose higher risk to epifauna are deposited. The most significant drawback to the method occurred during periods of low water clarity due to dense phytoplankton blooms. One important improvement would be to quantify the forces imparted by the lateral surge on individual BVL units. This could be accomplished with either velocimeter and OBS sensors which could be used to determine sediment flux, or with pressure transducers to measure force. Measuring fluxes across BVLs during a deployment would aid determining variation in the lateral extent of the deposition event, important when considering the overall or cumulative effect at a site.

V03. Example of benthic video lander deployment and comparison of normal, 5x, and 32x video speeds on behavioral observations of epifauna.

Sandy substrates of Pacific Northwest shallow nearshore zones are frequently disturbed by wave stress [27]. The largest impacts occur during winter storms but wave surges capable of mobilizing bottom sediments occur regularly [46]. Benthic fauna inhabiting these nearshore sites are adapted to disturbances by waves [15, 18], and as Video 2 demonstrates, we observed crabs to maneuver with dexterity in oscillatory flow strong enough for sediment resuspension. Organisms inhabiting sandy substrates also generally recover comparatively faster from disturbance than those at mud or silt substrates [19, 47, 48]. The energetic nearshore zone at the study site is naturally dispersive and so minimizes negative water quality impacts such as heightened turbidity [46, 49]. In fact, in many BVL trials the visibility increased in post-impact imagery as phytoplankton blooms were dispersed or flocculated by the lateral surge. The similarity in sediment grain size and composition between dredged and receiving areas [7] is another factor limiting negative effects of sediment deposition on the benthos [12].

Sediment deposition events from dredging activities have no natural analog at nearshore sites, except perhaps turbidity currents in submarine channels. Surge from storm events, both locally and remotely forced, generally build in intensity, allowing organisms to take precautionary actions such as burial or dispersal to deeper water. In contrast, the abrupt, forceful, and unanticipated impact of dredged sediment disposal plumes allows little opportunity for such behavioral adaptions, and we correspondingly observed crabs to be surprised and overwhelmed by the approaching lateral surge. Yet with the thin-layer disposal technique, the short duration of individual sedimentation events and limited deposition depths do not appear capable of inflicting acute harm to crab or whelk. What remains unclear is the experience of individual organisms entrained in the plume (including possible mechanical damage), as is whether the effects of multiple deposition events (cumulative impacts) has detrimental effects on epifauna. While chronic effects are under investigation [29], data to date suggests thin-layer disposal and the practice of staggering deposition tracks throughout the designated disposal area are warranted for reducing potential cumulative impacts on epifauna. Further observations targeting behavioral interactions would increase understanding of caveats regarding use of BRVs for sediment deposition studies, but results of this study indicate the methods described provide a standardized capability for assessing impacts to epifauna.

## Acknowledgements

We thank Captain Tim Stentz and crew of the *F/V Forerunner* and Brian Kelly and Jake Biron of Ocean Associates for their expertise during BVL deployments. Appreciation also goes to captains and crew of *Essayons* who ably guided the dredge through our moorings during deposition runs. Dale Beasley provided essential encouragement and guidance, and discussion with Hans R Moritz (USACE) and Kenneth Connell (NOAA) clarified aspects of sediment plume dynamics.

